# Targeting keratin 17-mediated reprogramming of *de novo* pyrimidine biosynthesis to overcome chemoresistance in pancreatic cancer

**DOI:** 10.1101/2022.08.24.504873

**Authors:** Chun-Hao Pan, Nina V. Chaika, Robert Tseng, Md Afjalus Siraj, Bo Chen, Katie L. Donnelly, Michael Horowitz, Cindy V. Leiton, Sumedha Chowdhury, Lucia Roa-Peña, Lyanne Oblein, Natalia Marchenko, Pankaj K. Singh, Kenneth R. Shroyer, Luisa F. Escobar-Hoyos

## Abstract

Pancreatic ductal adenocarcinoma (PDAC) is a leading cause of cancer death. We previously reported keratin 17 (K17) as a novel negative prognostic and predictive biomarker, whose overexpression confers the resistance to chemotherapies. Here, we investigated the mechanisms of chemoresistance and tumor-specific vulnerabilities that can be exploited for targeted therapies for K17-expressing PDAC. Unbiased metabolomic studies in isogenic PDAC models identified several key metabolic pathways that are upregulated in the presence of K17. We demonstrate that K17 increases pyrimidine biosynthesis, a pathway that has been linked to chemoresistance. Patient dataset analysis revealed that K17 expression and enzymes involved in pyrimidine, but not purine, *de novo* biosynthesis is associated with shorter patient survival. Rescue experiments showed that deoxycytidine (dC) and deoxythymidine (dT) were sufficient to promote resistance to Gemcitabine (a dC analog) and 5-fluorouracil (a dT analog), respectively. Furthermore, K17-expressing cells were more sensitive to Brequinar, a specific inhibitor of dihydroorotate dehydrogenase (DHODH), the rate-limiting enzyme in *de novo* pyrimidine biosynthetic pathway. Targeting DHODH by small interfering RNA or by Brequinar with Gemcitabine synergistically inhibited the viability of K17-positive PDAC cells. Importantly, the combination of Gemcitabine and Brequinar significantly inhibited the growth of K17-expressing tumors and extended survival of mice bearing K17-expressing PDACs. Overall, we identified a novel pathway of chemoresistance and a metabolic target of which could lead to the development of a biomarker-based therapy for K17-expressing PDAC.

## Introduction

Although the 5-year survival rate of pancreatic cancer has increased to 10% in 2020 in the United States^1^, it is still extremely low compared with other cancers. The most common form of pancreatic cancer, pancreatic ductal adenocarcinoma (PDAC) remains one of the deadliest cancers worldwide. Surgical resection is the only potentially curative intervention; however, approximately 80% of PDAC patients are not eligible for surgical procedure, making systemic therapeutic intervention their only option^2^. Unfortunately, PDAC is refractory to standard-of-care chemotherapy, responds to targeted therapies in less than 20% of cases that harbor actionable mutations, and does not respond to immunotherapy^2^. Thus, developing novel and more effective treatment strategies for PDAC is urgently needed to improve patient survival.

Compelling evidence has demonstrated that PDAC is comprised of molecular subtypes that are defined by gene expression signatures and correlate with differences in survival^3–6^. These findings highlight novel lines of investigation of surrogate biomarkers in molecular subtypes that may aid the development of more effective targeted therapy. We independently reported that keratin 17 (K17), one of the signature genes overexpressed in the Moffitt *basal-like* subtype^4^, is a novel negative prognostic and predictive biomarker in PDAC^7–9^. We also showed that K17 expression drives resistance to Gemcitabine (Gem) and 5-fluorouracil (5-FU)^9^, two key components of the main chemotherapeutic regimens for pancreatic cancer. Moreover, considering that there are no available therapeutic agents to inhibit K17, we performed a high-throughput drug screen and identified a small-molecule compound that selectively targets K17-expressing PDACs and increases survival of mice with K17-positive PADC^9^. In summary, our prior works have shown that K17 drives chemoresistance in PDAC and provides tumor-specific vulnerabilities that can be exploited for developing a biomarker-based therapy.

K17 is normally expressed during embryogenesis, silenced in mature somatic tissues except in certain stem cell populations^10,11^, and re-expressed in cancer^12,13^. Beyond its role as a cytoskeletal protein, K17 has been reported to impact the hallmarks of cancer by regulating key oncogenic signaling pathways^14^. Among these hallmarks^15^, metabolic reprogramming is one that has not been comprehensively evaluated. Research addressing altered cellular metabolism in cancer has been centered on understanding how metabolic pathway is rewired and whether targeting metabolic dependencies of cancer cells could be a selective anticancer strategy. In fact, some key cancer therapeutic regimens were developed by targeting specific metabolic pathways and remain effective in the clinic^16^. Here, we focused on studying whether K17 drives metabolic reprogramming and consequentially leads to chemoresistance. To address these questions, we performed unbiased metabolomic analyses and found that K17 reprograms several key metabolic pathways including up-regulating pyrimidine biosynthesis, which subsequently causes chemoresistance. This led us to identify that K17-expressing PDACs are more vulnerable to a selective inhibitor that targets *de novo* pyrimidine biosynthesis. In addition, targeting *de novo* pyrimidine biosynthesis enhances Gem-mediated cytotoxicity in K17-positive PDAC cells. In summary, we identified an unexpected pathway regulated by K17 that could ultimately lead to the development of a biomarker-based treatment for K17-exprssing PDAC.

## Materials and Methods

### Pathway and survival analyses from patient-derived data

Clinical annotations and transcripts per million RNA-seq data of TCGA PAAD samples^17^ were downloaded from UCSC xena data browser. Gene sets from Gene Ontology were downloaded from MSigDB collections. The GSVA R package was used to perform single-sample gene set enrichment analysis (ssGSEA) with gene expression of each sample screened against GMT gene set files, which transforms a gene-by-sample matrix into a pathway-by-sample matrix. The enrichment scores were normalized based on the ssGSEA method described in Barbie *et al*.^18^, which normalizes the scores by the absolute difference between minimum and maximum. K17 mRNA levels were correlated to normalized enrichment scores (NES) using the Spearman correlation test, and NES were compared between previously established high/low K17 groupings^9^ via Wilcoxon rank-sum test. Top metabolic pathways were chosen based on highest Spearman correlation, then significance.

### Cell culture

The human L3.6 PDAC cell line (Kras^G12A^)^19^ was kindly provided by Dr. Wei-Xing Zong (Rutgers University). PANC-1 PDAC cells (Kras^G12D^, p53R^273H^) were obtained from the American Type Culture Collection. The murine Kras^G12D^, p53^R172H^ pancreatic cancer (KPC) cell line was kindly provided by Dr. Gerardo Mackenzie (University of California at San Diego). Cells were cultured in a humidified incubator at 37°C under 5% CO_2_ in Dulbecco’s Modified Eagle Medium (DMEM, Gibco) supplemented with 10% fetal bovine serum (FBS, ThermoFisher) and 1% penicillin and streptomycin (P/S, Gibco).

### Genetic modification of K17 in PDAC cell lines

The generation of K17 gain-of-function cell line models (KPC and PANC-1) and K17 loss-of-function cell line model (L3.6) were previously described^9^. Briefly, to overexpress K17, cells were stably transduced to express either empty vector (EV) or human K17 (K17) in medium supplemented with 10% FBS for 24 hours, followed by fluorescence-activated cell sorting. Additionally, CRISPR-Cas9 mediated knockout technique was used to generate K17 loss-of-function cell line model.

### Protein extraction and Western blots

Cell lysates were prepared using RIPA buffer containing protease and phosphatase inhibitor cocktail (ThermoFisher Scientific). The lysates were sonicated, centrifuged, and the supernatant was collected. Protein concentration was then measured using the Bradford protein assay kit (Bio-Rad) according to the manufacturer’s instructions. Equal amounts of protein were separated by 10% SDS polyacrylamide gel electrophoresis. Immunoblotting was performed with primary antibodies to K17^20–22^ (a gift kindly provided by Dr. Pierre Coulombe, University of Michigan) and α-tubulin (Abcam), followed by infrared goat antimouse or goat anti-rabbit IgG secondary antibodies (LI-COR Inc.). Western blot images were captured by LI-COR Odyssey Imaging machine and images were quantified using Image Studio Lite software (LI-COR Inc.).

### Immunofluorescence imaging

As previously described^9^, cells were first fixed in ice-cold methanol at −20°C, permeabilized with 0.25% Triton X-100 at room temperature and blocked in PBS solution (Gibco) with 10% donkey serum (Sigma-Aldrich). Cells were incubated overnight with the primary K17 antibody^20^, diluted in 10% donkey serum. Fluorescein-conjugated goat anti-rabbit secondary antibody (Abcam) was incubated, and cells were counterstained with DAPI and mounted with Vectashield (Vector Laboratories).

### Mass spectrometric metabolomics analysis

Cells were plated in 6 cm plates and 2 hours before the collection of metabolites, the culture medium was replaced with fresh medium. Polar metabolites were extracted and then analyzed with LC-MS/MS using the selected reaction monitoring (SRM) method with positive/negative ion polarity switching on a Xevo TQ-S mass spectrometer^23,24^. Peak areas were integrated using MassLynx 4.1 (Waters Inc.), normalized to the respective protein concentrations, and the resultant peak areas were subjected to relative quantification analyses by utilizing Metaboanalyst 3.0 (www.metaboanalyst.ca)^25^.

### K17 immunohistochemistry

Immunohistochemical (IHC) staining and scoring were performed as previously described^7^. Briefly, slides were incubated at 60 °C, deparaffinized in xylene and rehydrated in alcohols. Antigen retrieval was performed using citrate buffer in a decloaking chamber at 120 °C for 10 minutes. To block endogenous peroxidase, 3% hydrogen peroxide was used. Tissue sections were incubated overnight at 4 °C with the rabbit monoclonal anti-K17 antibody (D12E5, Cell signaling, Danvers, MA). After incubation, biotinylated horse secondary antibodies (R.T.U. Vectastain ABC kit; Vector Laboratories, Burlingame, CA) were added. Development was done using 3, 3’ diaminobenzidine (DAB) (Dako, Carpinteria, CA) and counter-staining was done with hematoxylin. K17 staining intensity was classified by one pathologist (K.R.S) based on a subjective assessment of absent (0), light (+1), or strong (+2) staining. The overall proportion of tumor cells with 2+ intensity staining (the PathSQ score) was determined, blinded to corresponding data.

### Matrix assisted laser desorption/ionization mass spectrometry (MALDI MSI) metabolite imaging

Two K17 positive pancreatic tumor samples were collected from the KPC mouse model of PDAC for MALDI Imaging study. These tumors were prepared as snap frozen tissues immediately after harvesting. The freshly frozen tissues were sectioned on a Leica CM1510S Cryostat and were mounted on IntelliSlides. Prior to MSI analysis, tissue slides were dried in a vacuum desiccator at room temperature. 9-aminoacridine (9-AA) matrix solution was then sprayed onto the slide using an HTX M5 Sprayer. MALDI Imaging was performed on a timsTOF fleX mass spectrometer (Bruker Daltonics) over the entire section with a spatial resolution of 20 μm in negative ion mode. MSI images were generated, and the mean spectrum of MS signals was exported using SCiLS Lab software (Bruker Daltonics). Metabolites were annotated using MetaboScape software (Bruker Daltonics) with m/z tolerance set to 10 ppm and were cross validated with the Unified CCS compendium database. CMP level is normalized to the lowest expression baseline in each case. K17 IHC staining was performed on an adjacent serial section. The correlation of CMP and K17 was analyzed by capturing multiple voxels in a slide. Pearson’s correlation and *P*-value were calculated.

### Murine orthotopic xenograft studies

All experimental procedures were approved by the Institutional Animal Care and Use Committee (IACUC) at Stony Brook University and Yale University and are in accordance with the Guide for the Care and Use of Laboratory Animals from the National Institutes of Health. As previously described^9^, KPC cells with or without K17 expression were harvested during log-phase growth and resuspended in DMEM (Gibco) with Matrigel (Life Sciences) at a ratio of 1:1, to a final of 1,000 cells in a 30 μl volume. Cells were orthotopically implanted into the head of the pancreas of c57B6J mice. Tumor volume was tracked by ultrasound measurement. Once the tumor volume reached around 50 mm^3^, the mice were randomized into treatment groups and administered the following agents: Study I. Gem chemoresistance study-(1) vehicle or (2) Gem alone at 50 mg/kg body weight administered twice a week. Study II. 5-FU chemoresistance study-(1) vehicle or (2) 5-FU alone at 50 mg/kg body weight administered twice a week. Gem and 5-FU were given through intraperitoneal injections. Study III. Gem and Brequinar study-Immunodeficient NOD.SCID/Ncr mice were used for tumor implantation with human L3.6 K17 WT and K17 KO cells. Similarly, treatments started when tumors reached around 50 mm^3^. Gem at 25mg/kg was administered twice a week by intraperitoneal injection and Brequinar at 10 mg/kg was administered twice a week by gavage, for a total of 3 weeks. Tumor growth and body weight were monitored weekly. Survival was analyzed as an endpoint.

### Compounds tested

Gem (purity > 99%), 5-FU (purity > 99%), Brequinar (purity > 99%), Tetrahydrouridine (purity > 99%), 6-Mercaptopurine (purity > 99%), Azathioprine (purity > 99%), deoxyadenosine (purity > 99%), deoxyguanosine (purity > 99%), deoxycytidine (purity > 99%), and deoxythymidine (purity > 99%) were purchased from Sigma-Aldrich (St. Louis, MO, USA). The drugs were dissolved in 100% dimethyl sulfoxide (DMSO, Fisher BioReagents) with a stock concentration of 20 mM and were prepared for cell experiments at final DMSO concentrations at 0.1%. Nucleosides were dissolved in water and administered at 100 μM for drug response experiments.

### Cell viability and proliferation assays

Treatments of Gem and 5-FU and the supplement of deoxynucleosides were indicated in each experiment. Cell viability was examined using the CellTiter-Glo®Luminescent Cell Viability Assay (Promega), or the WST-1 Cell Proliferation Reagent (Sigma-Aldrich), using the method described previously^9,26^. Cells were plated and incubated overnight, drugs were added with 10-point dose-response titrations in triplicate (0.5 – 20,000 nM) or at the indicated concentration for 48 hours. To measure cell proliferation, cells were plated, and the relative proliferation index was measured by WST-1 assay at each time point.

### RNA isolation and qRT-PCR

Total RNA was extracted with TRIzol reagent (Life Technologies), according to the manufacturer’s instructions. cDNA synthesis was performed using MultiScribe™ Reverse Transcriptase (ThermoFisher Scientific), with 1 μg of RNA as a template for cDNA generation. TaqMan Universal Master Mix II, no UNG was used, and qRT-PCR was programmed using a QuantStudio 3 real-time PCR system (ThermoFisher Scientific). Data were normalized by the level of expression in each sample, as previously described^27^.

### Generation of DHODH knockdown cells

Transfection with the small interfering RNA (siRNA), was performed using the DharmaFECT™ system (Horizon Discovery Ltd., Cambridge, UK). Four constructs of siRNA: ON-TARGETplus siRNA J-009619-05 (target sequence: UAAAUUCCGAAAUCCAGUA), J-009619-06 (target sequence: CCACGGGAGAUGAGCGUUU), J-009619-07 (target sequence: GGACGGACUUUAUAAGAUG), and J-009619-08 (target sequence: GAACACAGGUUACGGGCCA); and Non-targeting siRNA (target sequence: UGGUUUACAUGUCGACUAA) were used. Briefly, cells were grown to 50% confluence in antibiotic-free 10% DMEM, then transfected with the plasmids for 24 hours according to the manufacturer’s protocol. Knockdown efficiency determined by Western blots was done 72 hours post-transfection. For the drug treatment experiment, cells were replaced with 10% DMEM with indicated drugs 24 hours post-transfection.

### Statistics

The half-maximal inhibitory concentration (IC50) of single-drug treatment was determined by GraphPad Prism 7 (Graph Pad Software). Drug pair interactions were calculated using the Highest Single Agent (HAS) model on the Combenefit software (Cancer Research UK Cambridge Institute, version 2.021)^28^. The statistical significance between the two groups was determined using Student’s *t*-test or one-way ANOVA test. The statistical significance between multiple variables was determined by two-way ANOVA test. Data were expressed as means ± standard deviation (SD) or standard error mean (SEM), and \**P*<0.05, \*\**P*<0.01, \*\*\**P*<0.001 were considered significant.

## Results

### K17 drives pyrimidine biosynthesis that is associated with short PDAC survival

Alterations in tumor metabolism are intimately linked to tumor aggression and chemoresistance^29–31^. K17 expression has been reported to impact cancer cell metabolism^14^; however, the detailed mechanisms remain unknown. To determine whether K17 promotes metabolic reprogramming, we analyzed primary PDAC tumor samples from The Cancer Genome Atlas (TCGA). We found that Biosynthesis, Metabolism, and Salvage pathways are significantly correlated to K17 mRNA expression and enriched in high-K17-expressing PDAC (**Fig. 1A**). Of note, pathways related to pyrimidine metabolism were those that had significantly higher enrichment scores. To further study how K17 impacts PDAC metabolism, we performed unbiased metabolomic studies of polar metabolites analysis in isogenic PDAC cell lines. We generated K17 gain-of-function (GOF) cell line models with murine KPC and human PANC-1 PDAC cell lines by stably transducing cells to express either empty vector (EV) or K17 (**Fig. 1B-C**). In addition, we used CRISPR-Cas9-mediated knockout method to generate K17 loss-of-function (LOF) cell line models with L3.6 human PDAC cells (**Fig. 1B-C**). We found that K17-expressing cells had significantly higher levels of intracellular purines and pyrimidines, including GTP, UTP, and CTP (**Fig. 1D-E**). In contrast, in K17 LOF models, the intracellular pools of purines and pyrimidines were overall reduced (**Fig. 1F**). Metabolites in pathways at the upstream of nucleotide metabolism, including glycolysis, tricarboxylic acid (TCA) cycle, and pentose phosphate pathway (PPP), were also significantly up-regulated in K17-expressing cells (**Fig. S1A-B**). Consistently, metabolic intermediates of glycolysis, TCA cycle, and PPP were lower in K17-negative cells (**Fig. S1C-D**). Specifically, in KPC K17-positive cells, high 6-phosphogluconate levels indicated that glucose was being routed through PPP to generate more nucleotides (**Fig. S1A**). High glutamine levels and a decrease in some TCA cycle metabolites with respect to succinate and 2-KG indicated the non-canonical pathway of glutamine metabolism that may facilitate nucleotide biosynthesis. Similar to KPC K17-expressing cells, in PANC-1 K17-positive cells, low levels of glyceraldehyde-3P and pyruvate coupled with a high level of 6-phosphogluconate indicated that glucose is being routed through PPP to generate more nucleotides. High glutamine and aspartate levels but low TCA cycle metabolites with respect to succinate, fumarate, and malate indicated that nucleotide biosynthesis was upregulated (**Fig. S1B**). This is contrary to the loss of K17 in L3.6 K17 knockout cells (**Fig. S1C**). To strengthen these findings, we used MALDI imaging to measure CMP levels in mouse tumor samples (**Fig. 1I**). Coupled with the corresponding K17 expression level (**Fig. 1J**), we found that K17 was significantly correlated with the level of CMP (**Fig. 1K**). Thus, these data in isogenic models suggest that independently of a host (mouse or human) and the model (GOF or LOF), K17 expression leads to metabolic reprogramming, resulting in increased intracellular pools of purines and pyrimidines.

**Figure 1:**
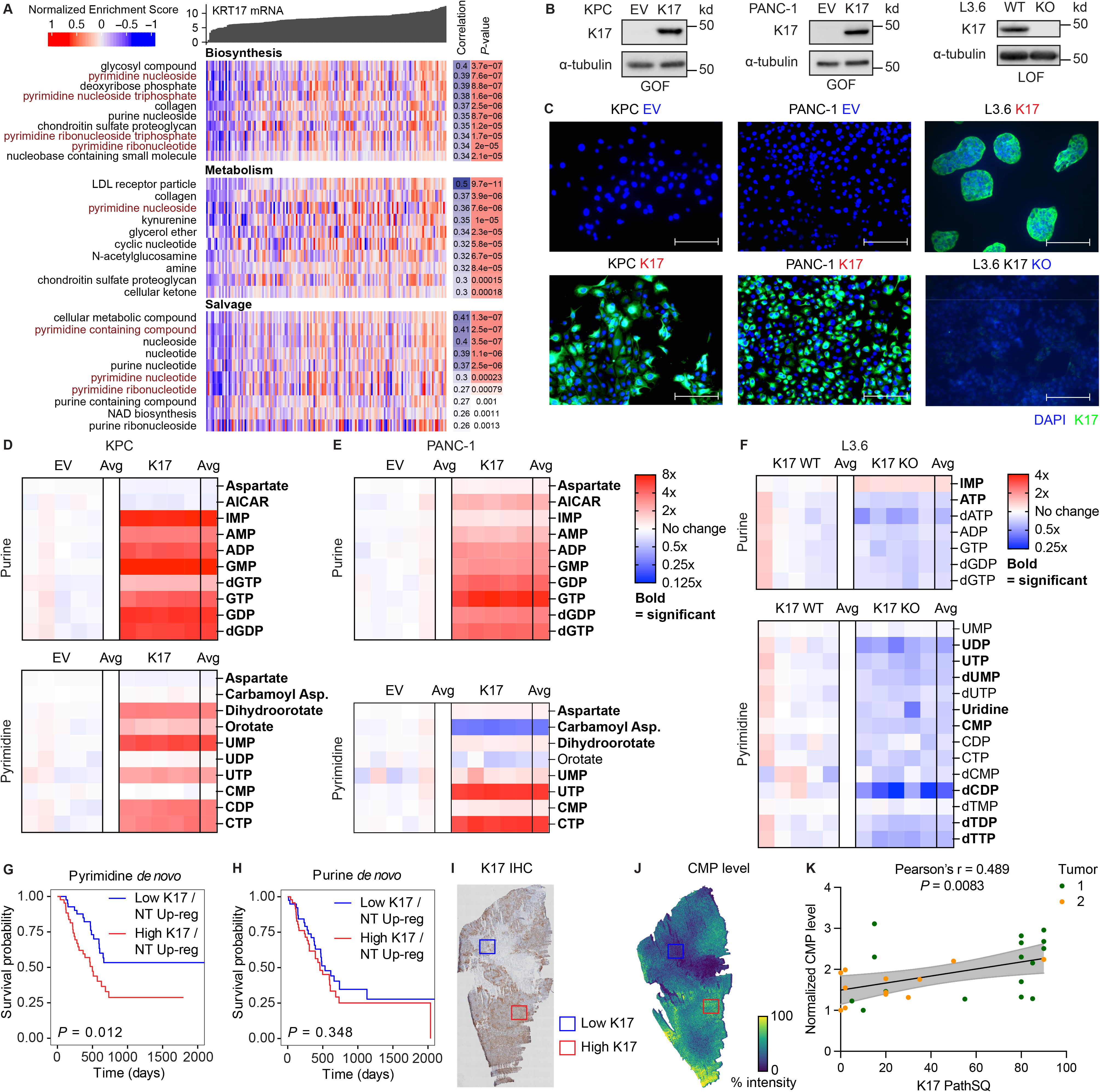
K17 drives enhanced nucleotide biosynthesis that is associated with short PDAC survival. **A.** K17 promotes metabolic reprogramming revealed by TCGA data analysis using primary PDAC tumor samples. Pyrimidine synthesis, purine synthesis, and salvage pathways are significantly correlated to K17 mRNA expression and enriched in high-K17-expressing PDAC. **B.** Genetic manipulation of K17 expression to generate stable cell line models. Murine Kras^G12D^, p53^R172H^ pancreatic cells (KPC) and human PANC-1 PDAC cells were transduced to stably express either empty vector (EV) or human K17 (K17). Bottom: CRISPR knockout of K17 (KO) and negative control (WT) in L3.6 human PDAC cell line to generate K17 loss-of-function (LOF) model. **C.** Immunofluorescence images of cell line models. Filament form of K17 is shown in green, and the nucleus is indicated by DAPI stain in blue. Scale bar = 100 μm. **D-F.** Metabolomic analyses in cell line models. K17 expression increased relative levels of purine and pyrimidine in nucleotide biosynthetic pathways (Student’s *t*-test, Bold=significant, n=5). **G-H.** Kaplan-Meier curves which stratify TCGA PDAC patients relative to KRT17 mRNA as well as their GSEA scores for nucleotide biosynthesis. High K17 cases with high up-regulated nucleotide metabolism in pyrimidine but not purine, were associated with shorten survival. **I.** Representative image K17 IHC staining. Two representative voxels highlight the area with low K17 (blue) and high K17 (red) expression. **J.** Representative image of CMP level measured by MALDI. The corresponding voxels with low- and high-K17 expression are highlighted. **K.** There is a significantly positive correlation of K17 expression with CMP level in mouse PDAC tissue.

We next studied how altered metabolic reprogramming of nucleotides and K17 expression impact patient survival. We analyzed TCGA datasets along with all pathways that were involved in nucleotide metabolism from the Gene Ontology database. We found that PDAC patients with up-regulated nucleotide metabolism (NM) had shorter survival, compared with those with down-regulated NM (data not shown). Based on our *in vitro* metabolomic findings that K17-expressing cells maintain higher intracellular pools of nucleotides, we next determined whether the up-regulation of these pathways involved in nucleotide biosynthesis along with high expression of K17 were associated with poor survival. Surprisingly, we found that up-regulated pyrimidine but not purine *de novo* biosynthesis was associated with short survival in high K17-expressing PDACs compared with low K17-expressing cases (log-rank *P* = 0.012 in pyrimidine vs. log-rank *P* = 0.348 in purine, **Fig. 1G-H**). K17 status did not further separate patient survival in nucleotide salvage pathways (log-rank *P* = 0.062 in pyrimidine vs. log-rank *P* = 0.778 in purine, **Fig. S1E-F**). These data suggest that even PDAC metabolism is rewired to promote tumor aggression, up-regulation of *de novo* pyrimidine biosynthesis and high K17 are key clinically-relevant factors that lead to short survival.

### Increased pools of deoxycytidine and deoxythymidine lead to resistance of Gem and 5-FU, respectively

Our previous work has demonstrated the role of K17 in driving chemoresistance to Gem and 5-FU, two major components in the PDAC first-line chemotherapeutic regimens^9^. In line with this, we showed that in animal models, treatment of Gem extended survival in mice bearing K17-negative tumors, while there was no survival difference in mice bearing K17-positive tumors after Gem treatments (log-rank *P* = 0.0001 in EV vs. log-rank *P* = 0.08 in K17, **Fig. 2A-B**). Interestingly, in our KPC animal models, treatment of 5-FU did not extend survival regardless of the expression of K17 (**Fig. S2A-B**). Notably, KPC tumors were highly resistant to 5-FU^9^. Of note, there was a trend that 5-FU treatment led to shorter survival in mice bearing K17-positive tumors compared with saline control (log-rank *P* = 0.14, **Fig. S2B**), suggesting that K17-expressing PDAC was extremely resistant to 5-FU treatment. While these studies reveal that K17-expressing PDAC is highly resistant to chemotherapeutic agents, the underlying mechanisms of chemoresistance have not been defined. Hence, we next determine whether K17 promotes chemoresistance by maintaining higher intracellular pools of nucleotides.

**Figure 2:**
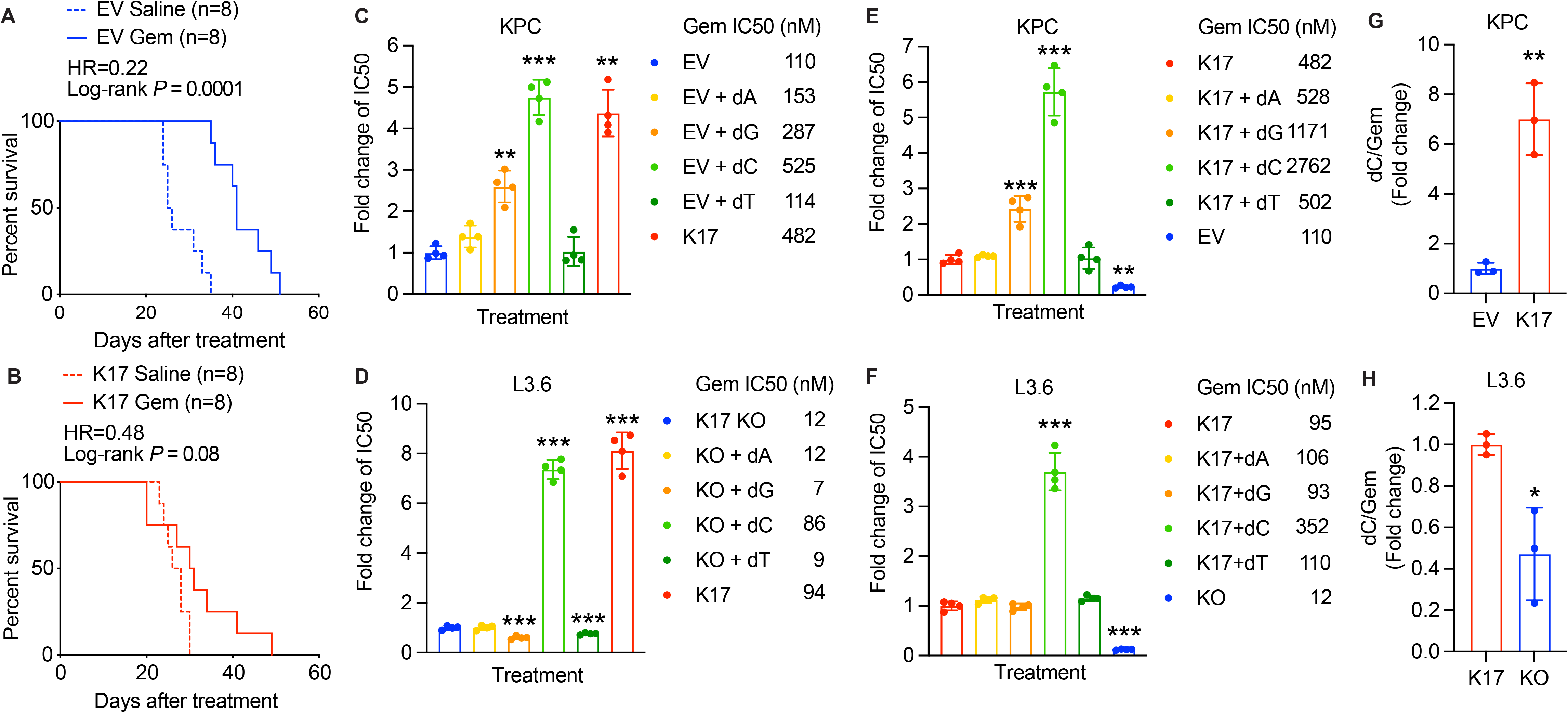
K17 mediates resistance to Gem by increasing intracellular deoxycytidine. **A-B.** Exogenous deoxycytidine (dC) increases Gem resistance in K17-negative (KPC EV and L3.6 KO) cells. KPC EV and L3.6 KO cells were treated with Gem combined with additionally individual nucleosides for 48h. The IC50 was determined by WST-1 assay followed by one-way ANOVA test. **C-D.** Exogenous deoxycytidine (dC) increases Gem resistance in K17-positive (KPC K17 and L3.6 K17 WT) cells. KPC K17 and L3.6 K17 cells were treated with Gem combined with additionally individual nucleosides for 48h. The IC50 was determined by WST-1 assay followed by one-way ANOVA test. **G-H.** K17-positive cells maintain higher dC/Gem ratio upon Gem treatment as determined by LC-MS/MS. Data were analyzed by one-way ANOVA test. \**P*<0.05, \*\**P*<0.01, \*\*\**P*<0.001

The increased pools of intracellular nucleotide have been reported to contribute to resistance to anticancer nucleoside analogs through the mechanism of molecular competition in PDAC^29,32^. Gem is a nucleoside analog, and when incorporated into DNA in place of cytosine, it prevents chain elongation and causes cell death^33^. Therefore, up-regulated nucleoside biosynthesis may impact its efficacy, as increased nucleotides could compete with Gem for DNA incorporation. Similar mechanism may exist in cells resistant to 5-FU, while 5-FU induce cell death with multiple mechanisms, including the inhibition of thymidylate synthase and the incorporation into RNA and DNA in place of uracil or thymine^34^. We next performed rescue experiments to study how individual nucleosides contribute to the resistance of Gem and 5-FU. We treated K17-negative PDAC cells with Gem and each nucleoside, deoxyadenosine (dA), deoxyguanosine (dG), deoxycytidine (dC), and deoxythymidine (dT), to assess which nucleoside(s) would lead to the highest resistance. We found that dC was sufficient to promote the highest resistance of Gem (a dC analog) in KPC K17-negative (EV) cells, compared with other nucleosides (**Fig. 2C**). Importantly, dC-driven resistance to Gem (IC50 = 525 nM) was at a comparable level of K17-positive cells only treated with Gem (IC50 = 482 nM). Similarly, in L3.6 K17 KO cells, the supplement of dC led to the highest resistance of Gem, compared with other nucleotides (**Fig. 2D**). We next evaluated the effect of nucleoside supplements on Gem sensitivity in K17-positive cells. As expected, supplements of dC induced the greatest effect of resistance to Gem in KPC and L3.6 K17-expressing cells (**Fig. 2E-F**). To validate that the dC-mediated resistance to Gem is a result of molecular competition, we performed LC-MS/MS metabolite analysis in our cell line models treated with Gem. We found that compared with K17-negative cells, K17-positive cells maintained a higher ratio of dC/Gem, suggesting that K17-expressing cells maintained higher intracellular dC levels over Gem (**Fig. 2G-H**). These observations are in line with other reports that increased level of dC contributes to Gem resistance^29,32^.

To determine if a similar molecular competition mechanism caused the resistance of 5-FU, we performed similar rescue experiments. In K17-negative cells (KPC EV and L3.6 K17 KO), dT drove the highest resistance of 5-FU (a dT analog), compared with other nucleosides (**Fig. 3A-B**). Interestingly, other nucleosides affected the sensitivity of 5-FU differently. This may be explained by some salvage mechanisms that cells can recycle ribose and nucleoside bases, which sequentially impacted the sensitivity. In K17-positive cells (KPC K17 and L3.6 K17), we found that only dT supplement led to or maintained the resistance to 5-FU (**Fig. 3C-D**, dark green bars). Of note, we did not observe significant differences in cell growth (**Fig. 3E-F**) or cell cycle progression (**Fig. S2C-E**) between PDAC cells with or without K17 expression. It rules out the possibility that increased pools of nucleotide was a reflect of rapid cell proliferation. As the uptake of nucleosides may also affect the efficacy of Gem and 5-FU, we also evaluated the expression of principle nucleoside transporters. The mRNA level of nucleoside transporters between K17-positive and K17-negative cells was similar (**Fig. 3G-H**), suggesting that the increased intracellular nucleosides may be a cell-autonomous mechanism. These data show for the first time that upregulation of pyrimidine biosynthesis by K17 underlies resistance to chemotherapeutic agents.

**Figure 3:**
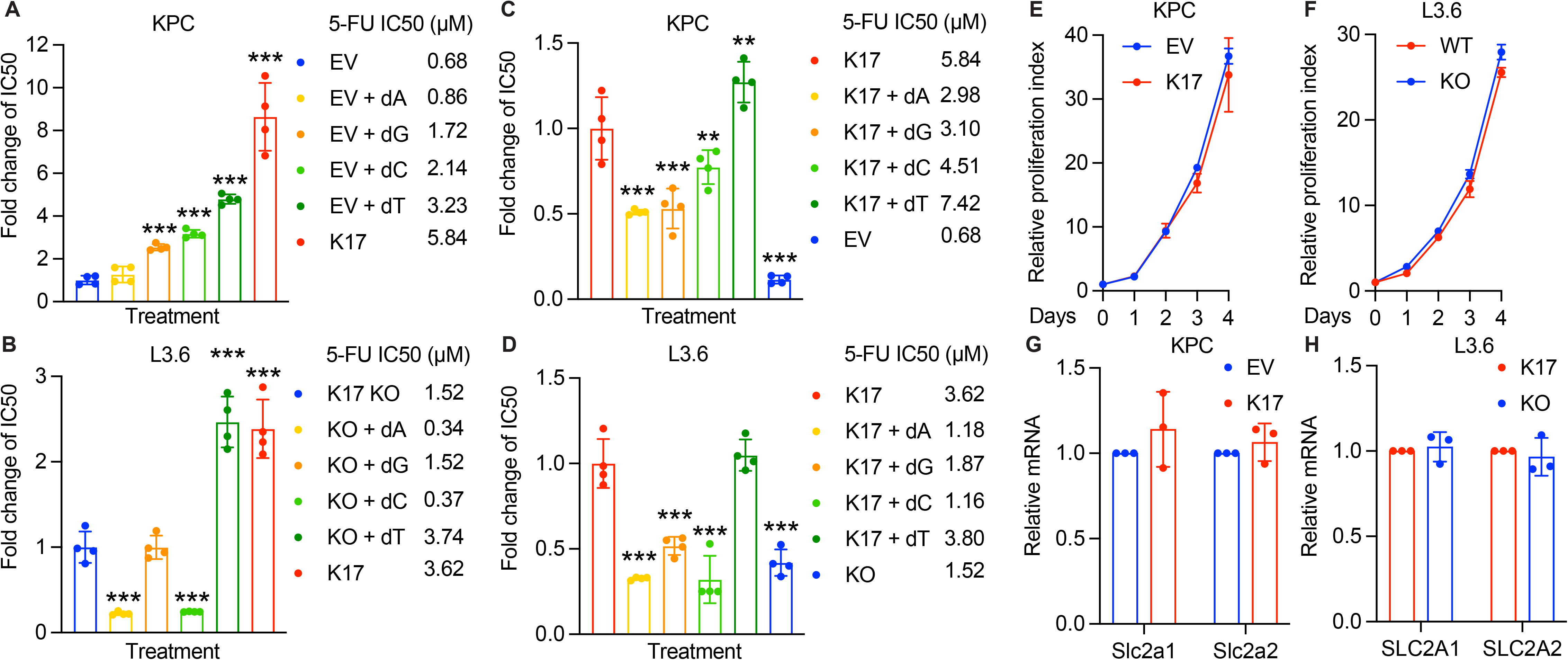
Increased pools of deoxythymidine drives resistance to 5-FU. **A-B.** Kaplan-Meier survival curves of mice bearing K17-negative and K17-positive tumors treated with 5-FU or saline control. N=8 per arm. **C-D.** Exogenous deoxythymidine (dT) increases 5-FU resistance in K17-negative (F-G) and K17-positive (H-I) cells. Cells were treated with 5-FU combined with additionally individual nucleosides for 48h. The IC50 was determined by WST-1 assay followed by one-way ANOVA test. **E-F.** No proliferation difference was found in KPC and L3.6 cell line models. **G-H.** No significant change at the mRNA level of enzymes involved in nucleoside transporter was found in KPC and L3.6 cell line models.

### K17-expressing PDAC cells show preferential sensitivity to inhibition of *de novo* pyrimidine biosynthesis

To determine whether targeting pyrimidine biosynthesis is a therapeutic opportunity in K17-expressing cells, we set out to test the nucleotide pathway dependency. We first reviewed literatures and came up with a list of nucleoside biosynthetic inhibitors that have been tested in cancer models^35,36^. Combined with the results from our previous high-throughput drug screen in K17 LOF cell line model^9^, we ruled out those that did not show sensitivity and specificity in K17-positive cells due to low response rates. Finally, we identified and tested four FDA-approved compounds that inhibit each nucleotide metabolic pathway. Using L3.6 cell line model, we found that compared with K17 WT cells, K17 KO cells were less sensitive to Brequinar, a compound that inhibits *de novo* pyrimidine biosynthesis (**Fig. 4A**). Brequinar has been tested and showed potent *in vitro* and *in vivo* anti-tumor activity across multiple tumor models^37,38^. In contrast, Tetrahydrouridine, a drug that inhibits pyrimidine salvage pathway, did not inhibit cell viability in L3.6 cells at a relatively higher dosage (**Fig. 4B**). Interestingly, L3.6 K17 WT cells were more resistant to both purine *de novo* and salvage inhibitors, Azathioprine and 6-Mercaptopurine (**Fig. 4C-D**). Coupled with our findings that up-regulation of pyrimidine but not purine *de novo* biosynthesis and high K17 expression in PDAC led to shorter survival, these data suggest that K17 expression promotes nucleotide biosynthesis, and K17-expressing PDAC cells depend on *de novo* pyrimidine biosynthetic pathway, presenting as a novel therapeutic target.

**Figure 4:**
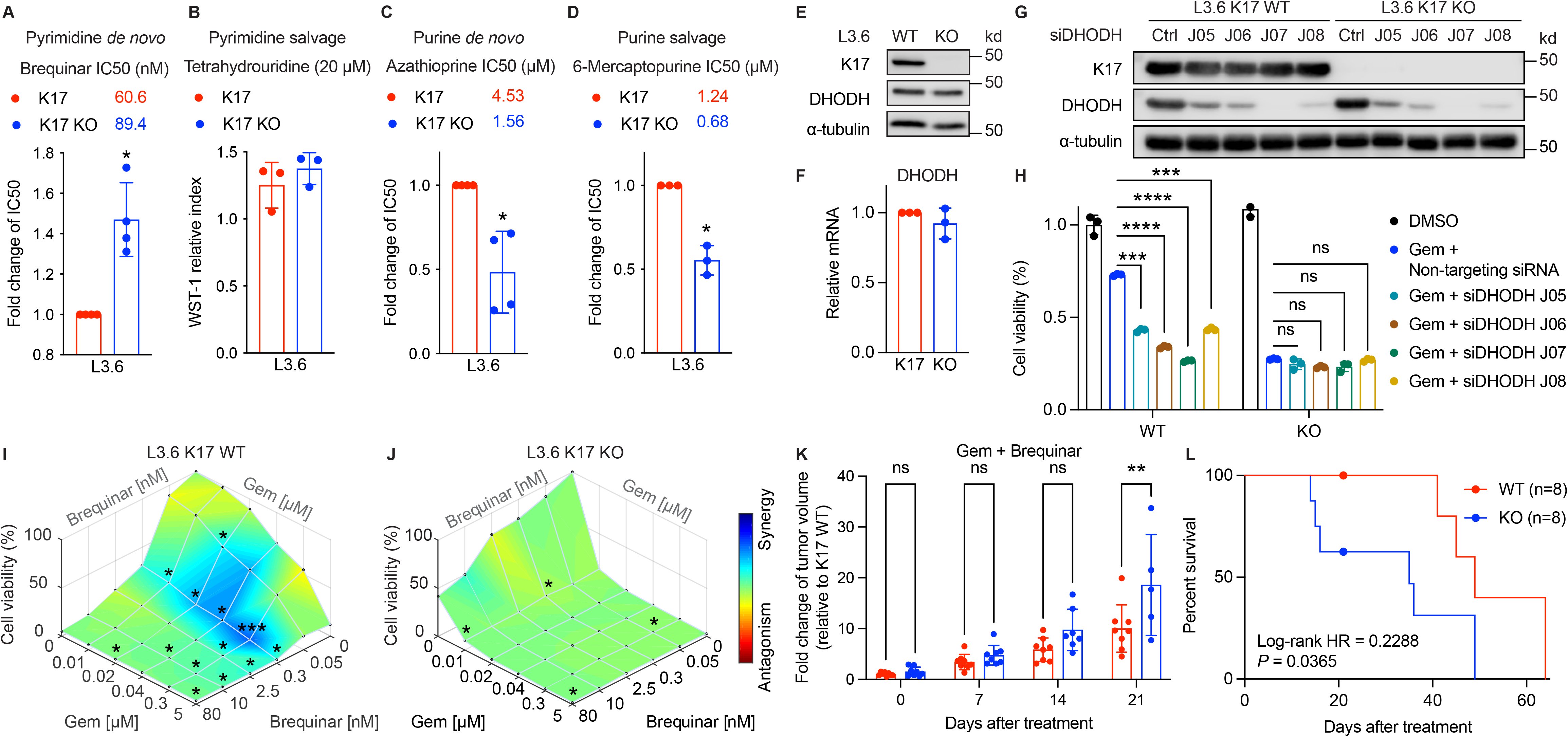
Targeting pyrimidine *de novo* biosynthesis is a therapeutic opportunity in K17-expressing PDAC. **A.** L3.6 cells expressing K17 are more sensitive to Brequinar, a drug that inhibits pyrimidine *de novo* biosynthesis, with significantly lower IC50 value. **B.** Tetrahydrouridine, a drug that inhibits pyrimidine salvage biosynthesis, did not inhibit cell viability in L3.6 cells. **C-D.** L3.6 cells expressing K17 are more resistant to purine *de novo* and salvage inhibitors. **E-F.** The protein (E) and mRNA (F) levels of DHODH in L3.6 K17 LOF cell line models. **G.** Western blots show that siRNA against DHODH knocked down the expression of DHODH in L3.6 K17 WT and K17 KO cells. **H.** The cell viability of L3.6 K17 WT and K17 KO cells transfected with siRNAs against DHODH or non-targeting siRNA with the treatment of Gem at 500 nM. Two-way ANOVA test. **I-J.** Combination treatment of Brequinar and Gem in K17-positive (K) and K17-negative (L) cells. The dose response surface was shown with the synergy distribution indicated by colors. **K-L.** The combination of Gem and Brequinar shows higher anticancer effects in K17-expressing PDACs. With the combination treatment, tumor volumes of L3.6 K17-expressing tumors (K17 WT) were significantly decreased, compared with L3.6 tumors with K17 KO (K). The reatment of Gem and Brequinar significantly extended the survival of animals bearing K17-expressing (K17 WT) tumors. \**P*<0.05, \*\*\**P*<0.001, \*\*\*\**P*<0.0001

Given the fact that Brequinar is selectively toxic in K17-positive PDAC cells, we analyzed the expression of its target, dihydroorotate dehydrogenase (DHODH, the rate-limiting enzyme in *de novo* pyrimidine biosynthesis) with regard to K17 expression. We did not observe significant differences in DHODH expression between L3.6 K17 WT and K17 KO cells at protein and mRNA levels (**Fig. 4E-F**). Next, we performed experiments to evaluate whether the knockdown of DHODH sensitized K17-expressing cells to Gem. Using four different sequence constructs of siRNA against DHODH and non-targeting siRNA control, we generated four DHODH LOF cell line models in L3.6 K17 WT cells (**Fig. 4G**). We then tested Gem sensitivity in these K17-positive cells with altered DHODH expression. We found that compared with non-targeting control, knockdown of DHODH led to lower cell viability upon Gem treatment (**Fig. 4H**), suggesting that K17-expressing cells became sensitive to Gem with the interruption at the *de novo* pyrimidine biosynthesis. Importantly, in K17-negative cells, we did not observe such effects by knocking down DHODH followed by Gem treatment (**Fig. 4G-H**). These data suggest that K17-expressing PDAC cells depend on *de novo* pyrimidine biosynthesis, and DHODH is a critical enzyme that contributes resistance to Gem.

Lastly, we validated these findings using Brequinar and Gem. To investigate whether Brequinar would enhance the Gem-mediated cytotoxic effect and determine whether the combined effect is synergistic or antagonistic, we performed co-treatment experiments. We tested Brequinar and Gem with five increasing doses, and the combination of Brequinar and Gem in the five times five dosing format. The cell viability under combinatorial treatments was analyzed by Combenefit^28^, a software that performs synergy analyses. As expected, L3.6 K17 WT cells (**Fig. 4I**) were more resistant to the single treatment of Gem compared with L3.6 K17 KO cells (**Fig. 4J**), as indicated by the dose-response cell viability curve against the z-axis on the right end side of the graph. For Brequinar, K17 WT cells were more sensitive than K17 KO cells as shown in the left end side of the graph. For combination treatments, the dose-response surface was shown with the synergy distribution indicated by colors. The combination of Gem and Brequinar showed synergistic effects in K17 WT cells, while there were neutral effects of the combination in K17 KO cells. We also observed that by treating cells with a low dose of Brequinar, the cell viability was significantly lower than Gem single treatment along with in K17-positive cells. To strength our *in vitro* findings, we performed an orthotopic xenograft animal study and evaluated the anticancer effects of Gem and Brequinar. At day 21 post-combination treatment, the combination of Gem plus Brequinar significantly inhibited the growth of K17-expressing tumors (**Fig. 4K**, tumor sizes of K17 WT tumors were half of the size of K17 KO tumors). In addition, the combination of Gem and Brequinar significantly extended survival of mice bearing K17 WT tumors (median survival of 49 days) compared with mice with K17 KO tumors (median survival of 35 days) (**Fig. 4L**, HR = 0.2288, Log-rank *P* = 0.0365). These data suggest that Brequinar is a potent drug to target K17-expressing PDAC cells, and when combined with Gem, the two drugs work synergistically for K17-expressing PDAC.

## Discussion

PDAC is a malignancy with very poor prognosis, and limited progress has been made on the development of novel treatment strategies that can improve therapeutic efficacy. The intrinsic resistance to chemotherapy is one of the major reasons that PDAC patients have relatively short survival^39,40^. We previously reported that K17, a novel biomarker of the most lethal molecular subtype of PDAC, can be a powerful tool to predict patient survival and response to chemotherapy^7–9^. K17 also promotes resistance to Gem and 5-FU, while the molecular mechanisms that drive chemoresistance remain undefined. To develop novel and potentially more effective biomarker-based therapies for PDAC, we aimed to investigate the mechanism of K17-induced chemoresistance and determine if targeting K17-dependent pathways would enhance therapeutic response.

Certain chemoresistant mechanisms of anti-cancer nucleoside analogs have been depicted, relating to drug transporters, the activation or inactivation of metabolic enzymes, and the tumor microenvironment^40^. For example, the resistance of Gem has been reported to be caused by limited uptake of Gem due to reduced nucleoside transporters^41^, and up-regulated dC biosynthesis^29^ or increased dC consumption from the tumor microenvironment that directly competes and reduces its effectiveness^32^. On the other hand, multiple factors have been suggested to contribute to 5-FU resistance, including increased expression of thymidylate synthase, increased activity of deoxyuridine triphosphatase, and overexpression of anti-apoptotic proteins^34^. However, how K17 drives resistance to Gem and 5-FU remains largely unclear. Here, we focused on studying whether altered metabolism is a key contributor of K17-induced chemoresistance.

There are limited reports summarizing how K17 impacts tumor metabolism. A study has shown that K17 impacts cancer cell metabolism by the Akt/mTOR/hypoxia-inducible factor (HIF)-1α pathway in osteosarcoma, and the knockdown of K17 inhibited glycolysis *in vitro*^42^. However, whether K17 drives chemoresistance by metabolic rewiring has not been studied in PDAC. To explore how K17 impacts metabolism, we performed unbiased *in silico* and *in vitro* metabolomic analyses and found that K17 expression up-regulates several key metabolic pathways and contributes to chemoresistance. These findings are novel as they connect an oncoprotein to critical pathways of nucleotide metabolism, especially *de novo* pyrimidine biosynthesis. It also provides a rationale to explore the therapeutic opportunity by targeting K17-induced pyrimidine biosynthesis.

In addition, the finding that increased intracellular pools of dC lead to resistance to Gem validates a similar molecular competition mechanism of chemoresistance from previous publications^29,32^. These data strengthen the fact that altered metabolism is one of a major driving force of chemoresistance. In our case, increased pools of dC in PDAC cells led to decreased uptake and sensitivity to Gem. Interestingly, we found a similar trend of molecular competition between dT and 5-FU, which has not been reported previously. Of note, the rescue experiment with 5-FU and other nucleosides, including dA, dG, and dC, made cells more sensitive to 5-FU. As known, 5-FU has a much more complicated manner in terms of the mechanism of actions to induce cell death^43^. It has also been reported that the imbalances of deoxynucleotides would severely disrupt DNA synthesis and repair, resulting in lethal DNA damage and causing cell death^44^. This may explain why under 5-FU treatment, the supplement of dA, dG, and dC lead to more cell death due to the imbalanced pools of deoxynucleotides, while future research remains to be done.

Our patient data analyses showed that the upregulation of pyrimidine but not purine *de novo* biosynthesis is associated with short survival when PDACs express a high level of K17. In addition, we evaluated that K17-expressing cells are more vulnerable to an inhibitor that interrupts *de novo* pyrimidine biosynthesis and resistant to purine biosynthesis inhibitors. These data show for the first time the metabolic dependency of K17-positive tumors on pyrimidine but not purine, and a potential therapeutic opportunity by targeting K17-dependent metabolic pathway. This unique dependency may reform the rationale of treatment strategy design to focus on targeting pyrimidine *de novo* biosynthesis in PDAC. Interestingly, the inhibitor targeting pyrimidine salvage was not effective in killing K17-positive cells. This points out that K17-expressing cells may rely more on the *de novo* than salvage pathway to synthesize pyrimidine.

Using genetic knockdown or pharmacologic treatment to inhibit DHODH, the rate-limiting enzyme for *de novo* pyrimidine biosynthesis, we were able to make K17-expressing cells re-sensitized to Gem. Of note, siRNA knockdown of DHODH did not significantly impact cell viability of K17-expressing cells. This may be because pharmacologic treatment of Brequinar to directly inhibit DHODH induces cell death more efficiently than genetic alteration, or cells transfected with siRNAs against DHODH may compensate nucleotide production more easily. To this end, how K17 impacts DHODH is still unexplored. Our data suggest that there is no differentially expressed change of DHODH at protein and mRNA levels between cells with or without K17. Thus, the mechanistic connection between K17 and DHODH may beyond the regulation of transcription and translation. As DHODH is a mitochondrial enzyme, how K17 may impact mitochondria and DHODH, and subsequentially upregulate *de novo* pyrimidine biosynthesis would be worth investigating.

Finally, we demonstrated that Brequinar works synergistically with Gem in K17-expressing PDAC cells. Encouragingly, our animal model showed that combination therapy significantly inhibited the tumor growth of K17-positive PDACs and extended the survival of mice bearing K17-expressing tumors. While Brequinar has previously been found to have promising *in vitro* and *in vivo* anti-cancer activity in multiple tumor models, including melanoma^45^, lung cancer^46^, and pancreatic cancer^47^, it was not found in the 1990s to provide significant, objective responses in clinical trials of patients with solid tumors^48–50^. The failure of Brequinar to impact tumor growth in these early studies might have been undermined by the absence of predictive biomarkers at that time to guide clinical enrollment. Here, we demonstrate that K17 is a powerful biomarker to identify tumors that may benefit the most from Brequinar treatment. Future studies could further explore whether more selective and potent compounds that target DHODH can be designed for treating K17-expressing PDACs.

### Conclusions

We define a novel connection between K17, nucleotide metabolism, and chemoresistance in PDAC. We report for the first time that K17, a negative biomarker of the most aggressive PDAC subtype, drives chemoresistance by up-regulating *de novo* pyrimidine biosynthesis, which provides a tumor-specific vulnerability: DHODH. Although future studies are needed to determine why K17-positive PDAC depends on *de novo* pyrimidine biosynthesis, we have already provided evidence to support the development of potential metabolic targeted therapies for K17-expressing PDAC.

## Acknowledgements

The authors thank the Biological Mass Spectrometry Shared Resource at the Stony Brook Cancer Center for assisting the MALDI-MSI analysis and the Yale Center for Molecular Discovery for conducting critical experimets. We also appriciate the help from members of our laboratories for critical reading of the manuscript.

## Legends

**Supplementary Figure 1: Metabolomic analysis in K17-manipulated cell line models and patient-derived survival analysis.**

**A-C.** Metabolomic analyses in cell line models. K17 expression increased relative levels of glycolysis, TCA cycle, and PPP (Student’s *t*-test, Bold=significant, n=5).

**D.** Kaplan-Meier curves separated by up-regulated or down-regulated nucleotide metabolism (NM) and low K17 or high K17 status.

**E-F.** Kaplan-Meier curves of patients with up-regulated pyrimidine (E) or purine (F) salvage NM, further separated by K17 status.

**Supplementary Figure 2: Intracellular nucleosides impact sensitivity to Gem and 5-FU. A-B.** Kaplan-Meier survival curves of mice bearing K17-negative and K17-positive tumors treated with 5-FU or saline control. N=8 per arm.

**C-E.** No difference was found in cell cycle progression between K17-positive and K17-negative cells in tested cell line models. \**P*<0.05, \*\**P*<0.01, \*\*\**P*<0.001

## LITERATURE CITED

1 Siegel, R. L., Miller, K. D., Fuchs, H. E. & Jemal, A. Cancer Statistics, 2021. CA Cancer J Clin 71, 7–33 (2021). https://doi.org:10.3322/caac.21654

2 Kleeff, J. et al. Pancreatic cancer. Nat Rev Dis Primers 2, 16022 (2016). https://doi.org:10.1038/nrdp.2016.22

3 Collisson, E. A. et al. Subtypes of pancreatic ductal adenocarcinoma and their differing responses to therapy. Nat Med 17, 500–503 (2011). https://doi.org:10.1038/nm.2344

4 Moffitt, R. A. et al. Virtual microdissection identifies distinct tumor-and stroma-specific subtypes of pancreatic ductal adenocarcinoma. Nat Genet 47, 1168–1178 (2015). https://doi.org:10.1038/ng.3398

5 Bailey, P. et al. Genomic analyses identify molecular subtypes of pancreatic cancer. Nature 531, 47–52 (2016). https://doi.org:10.1038/nature16965

6 Collisson, E. A., Bailey, P., Chang, D. K. & Biankin, A. V. Molecular subtypes of pancreatic cancer. Nat Rev Gastroenterol Hepatol 16, 207–220 (2019). https://doi.org:10.1038/s41575-019-0109-y

7 Roa-Peña L et al. Keratin 17 identifies the most lethal molecular subtype of pancreatic cancer. Scientific reports 9 (2019).

8 Roa-Pena, L. et al. Keratin 17 testing in pancreatic cancer needle aspiration biopsies predicts survival. Cancer Cytopathol (2021). https://doi.org:10.1002/cncy.22438

9 Pan, C. H. et al. An unbiased high-throughput drug screen reveals a potential therapeutic vulnerability in the most lethal molecular subtype of pancreatic cancer. Molecular Oncology 14, 1800–1816 (2020). https://doi.org:10.1002/1878-0261.12743

10 Moll, R., Franke, W. W., Schiller, D. L., Geiger, B. & Krepler, R. The catalog of human cytokeratins: patterns of expression in normal epithelia, tumors and cultured cells. Cell 31, 11–24 (1982).

11 Chu, P. G. & Weiss, L. M. Keratin expression in human tissues and neoplasms. Histopathology 40, 403–439 (2002). https://doi.org:https://doi.org/10.1046/j.1365-2559.2002.01387.x

12 Rhodes, D. R. et al. ONCOMINE: A Cancer Microarray Database and Integrated Data-Mining Platform. Neoplasia 6, 1–6 (2004). https://doi.org:https://doi.org/10.1016/S1476-5586(04)80047-2

13 Babu, S. et al. Keratin 17 is a sensitive and specific biomarker of urothelial neoplasia. Modern Pathology 32, 717–724 (2019). https://doi.org:10.1038/s41379-018-0177-5

14 Baraks, G. et al. Dissecting the Oncogenic Roles of Keratin 17 in the Hallmarks of Cancer. Cancer Res (2021). https://doi.org:10.1158/0008-5472.CAN-21-2522

15 Hanahan, D. Hallmarks of Cancer: New Dimensions. Cancer Discov 12, 31–46 (2022). https://doi.org:10.1158/2159-8290.CD-21-1059

16 Vander Heiden, M. G. Targeting cancer metabolism: a therapeutic window opens. Nat Rev Drug Discov 10, 671–684 (2011). https://doi.org:10.1038/nrd3504

17 Nawy, T. A pan-cancer atlas. Nature Methods 15, 407–407 (2018). https://doi.org:10.1038/s41592-018-0020-4

18 Barbie, D. A. et al. Systematic RNA interference reveals that oncogenic KRAS-driven cancers require TBK1. Nature 462, 108–112 (2009). https://doi.org:10.1038/nature08460

19 Bruns, C. J., Harbison, M. T., Kuniyasu, H., Eue, I. & Fidler, I. J. In vivo selection and characterization of metastatic variants from human pancreatic adenocarcinoma by using orthotopic implantation in nude mice. Neoplasia 1, 50–62 (1999). https://doi.org:10.1038/sj.neo.7900005

20 McGowan, K. M. & Coulombe, P. A. Onset of keratin 17 expression coincides with the definition of major epithelial lineages during skin development. J Cell Biol 143, 469–486 (1998). https://doi.org:10.1083/jcb.143.2.469

21 Depianto, D., Kerns, M. L., Dlugosz, A. A. & Coulombe, P. A. Keratin 17 promotes epithelial proliferation and tumor growth by polarizing the immune response in skin. Nat Genet 42, 910–914 (2010). https://doi.org:10.1038/ng.665

22 Hobbs, R. P. et al. Keratin-dependent regulation of Aire and gene expression in skin tumor keratinocytes. Nat Genet 47, 933–938 (2015). https://doi.org:10.1038/ng.3355

23 Gunda, V., Yu, F. & Singh, P. K. Validation of Metabolic Alterations in Microscale Cell Culture Lysates Using Hydrophilic Interaction Liquid Chromatography (HILIC)-Tandem Mass Spectrometry-Based Metabolomics. PLoS One 11, e0154416 (2016). https://doi.org:10.1371/journal.pone.0154416

24 Yuan, M., Breitkopf, S. B., Yang, X. & Asara, J. M. A positive/negative ion-switching, targeted mass spectrometry-based metabolomics platform for bodily fluids, cells, and fresh and fixed tissue. Nat Protoc 7, 872–881 (2012). https://doi.org:10.1038/nprot.2012.024

25 Xia, J. & Wishart, D. S. Using MetaboAnalyst 3.0 for Comprehensive Metabolomics Data Analysis. Curr Protoc Bioinformatics 55, 14 10 11–14 10 91 (2016). https://doi.org:10.1002/cpbi.11

26 Pan, C. H. et al. Vorinostat enhances the cisplatin-mediated anticancer effects in small cell lung cancer cells. BMC Cancer 16, 857 (2016). https://doi.org:10.1186/s12885-016-2888-7

27 Schmittgen, T. D. & Livak, K. J. Analyzing real-time PCR data by the comparative C(T) method. Nat Protoc 3, 1101–1108 (2008). https://doi.org:10.1038/nprot.2008.73

28 Di Veroli, G. Y. et al. Combenefit: an interactive platform for the analysis and visualization of drug combinations. Bioinformatics 32, 2866–2868 (2016). https://doi.org:10.1093/bioinformatics/btw230

29 Shukla, S. K. et al. MUC1 and HIF-1alpha Signaling Crosstalk Induces Anabolic Glucose Metabolism to Impart Gemcitabine Resistance to Pancreatic Cancer. Cancer Cell 32, 71–87 e77 (2017). https://doi.org:10.1016/i.ccell.2017.06.004

30 Goode, G. et al. MUC1 facilitates metabolomic reprogramming in triple-negative breast cancer. PLoS One 12, e0176820 (2017). https://doi.org:10.1371/journal.pone.0176820

31 Hattori, A. et al. Cancer progression by reprogrammed BCAA metabolism in myeloid leukaemia. Nature 545, 500–504 (2017). https://doi.org:10.1038/nature22314

32 Halbrook, C. J. et al. Macrophage-Released Pyrimidines Inhibit Gemcitabine Therapy in Pancreatic Cancer. Cell Metab 29, 1390–1399 e1396 (2019). https://doi.org:10.1016/j.cmet.2019.02.001

33 Plunkett, W. et al. Gemcitabine: metabolism, mechanisms of action, and self-potentiation. Semin Oncol 22, 3–10 (1995).

34 Zhang, N., Yin, Y., Xu, S. J. & Chen, W. S. 5-Fluorouracil: mechanisms of resistance and reversal strategies. Molecules 13, 1551–1569 (2008). https://doi.org:10.3390/molecules13081551

35 Mathews, C. K. Deoxyribonucleotide metabolism, mutagenesis and cancer. Nat Rev Cancer 15, 528–539 (2015). https://doi.org:10.1038/nrc3981

36 Okesli, A., Khosla, C. & Bassik, M. C. Human pyrimidine nucleotide biosynthesis as a target for antiviral chemotherapy. Curr Opin Biotechnol 48, 127–134 (2017). https://doi.org:10.1016/j.copbio.2017.03.010

37 Sykes, D. B. The emergence of dihydroorotate dehydrogenase (DHODH) as a therapeutic target in acute myeloid leukemia. Expert Opin Ther Targets 22, 893–898 (2018). https://doi.org:10.1080/14728222.2018.1536748

38 Zhou, Y. et al. DHODH and cancer: promising prospects to be explored. Cancer Metab 9, 22 (2021). https://doi.org:10.1186/s40170-021-00250-z

39 Chand, S., O’Hayer, K., Blanco, F. F., Winter, J. M. & Brody, J. R. The Landscape of Pancreatic Cancer Therapeutic Resistance Mechanisms. Int J Biol Sci 12, 273–282 (2016). https://doi.org:10.7150/ijbs.14951

40 Zeng, S. et al. Chemoresistance in Pancreatic Cancer. Int J Mol Sci 20 (2019). https://doi.org:10.3390/ijms20184504

41 Rauchwerger, D. R., Firby, P. S., Hedley, D. W. & Moore, M. J. Equilibrative-sensitive nucleoside transporter and its role in gemcitabine sensitivity. Cancer Res 60, 6075–6079 (2000).

42 Yan, X. et al. Knockdown of KRT17 decreases osteosarcoma cell proliferation and the Warburg effect via the AKT/mTOR/HIF1alpha pathway. Oncol Rep 44, 103–114 (2020). https://doi.org:10.3892/or.2020.7611

43 Longley, D. B., Harkin, D. P. & Johnston, P. G. 5-fluorouracil: mechanisms of action and clinical strategies. Nat Rev Cancer 3, 330–338 (2003). https://doi.org:10.1038/nrc1074

44 Jarmula, A., Dowiercial, A. & Rode, W. A molecular modeling study of the interaction of 2’-fluoro-substituted analogues of dUMP/FdUMP with thymidylate synthase. Bioorg Med Chem Lett 18, 2701–2708 (2008). https://doi.org:10.1016/j.bmcl.2008.03.016

45 Dorasamy, M. S., Ab, A., Nellore, K. & Wong, P. F. Synergistic inhibition of melanoma xenografts by Brequinar sodium and Doxorubicin. Biomed Pharmacother 110, 29–36 (2019). https://doi.org:10.1016/j.biopha.2018.11.010

46 Li, L. et al. Identification of DHODH as a therapeutic target in small cell lung cancer. Sci Transl Med 11 (2019). https://doi.org:10.1126/scitranslmed.aaw7852

47 Santana-Codina, N. et al. Oncogenic KRAS supports pancreatic cancer through regulation of nucleotide synthesis. Nat Commun 9, 4945 (2018). https://doi.org:10.1038/s41467-018-07472-8

48 Dodion, P. F. et al. Phase II trial with Brequinar (DUP-785, NSC 368390) in patients with metastatic colorectal cancer: a study of the Early Clinical Trials Group of the EORTC. Ann Oncol 1, 79–80 (1990). https://doi.org:10.1093/oxfordjournals.annonc.a057680

49 Moore, M. et al. Multicenter phase II study of brequinar sodium in patients with advanced gastrointestinal cancer. Invest New Drugs 11, 61–65 (1993). https://doi.org:10.1007/BF00873913

50 Maroun, J. et al. Multicenter phase II study of brequinar sodium in patients with advanced lung cancer. Cancer Chemother Pharmacol 32, 64–66 (1993). https://doi.org:10.1007/BF00685878

