## Supplementary figures and images for "Targeting keratin 17-mediated reprogramming of *de novo* pyrimidine biosynthesis to overcome chemoresistance in pancreatic cancer"

### Figure S1

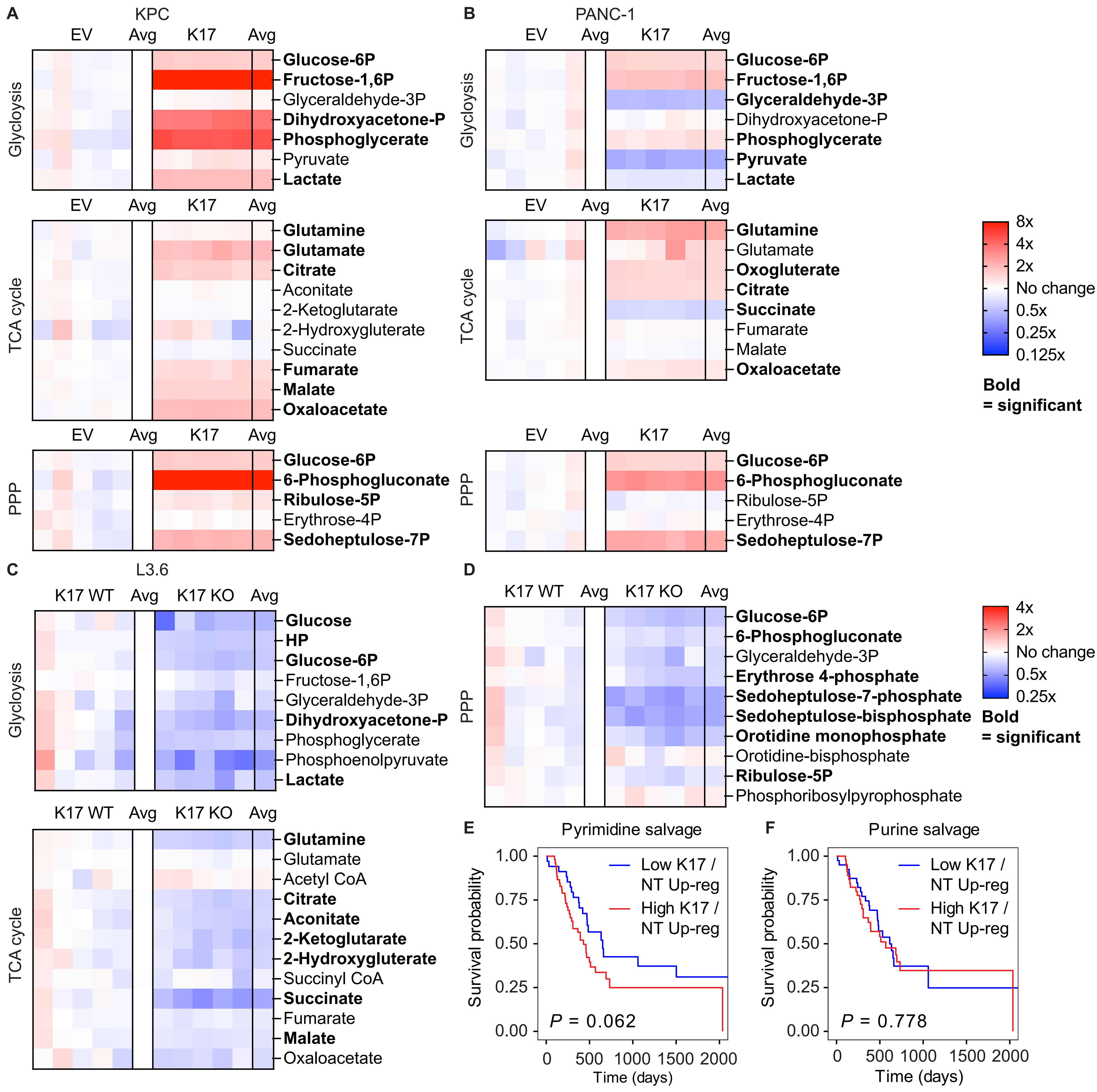

### Figure S2

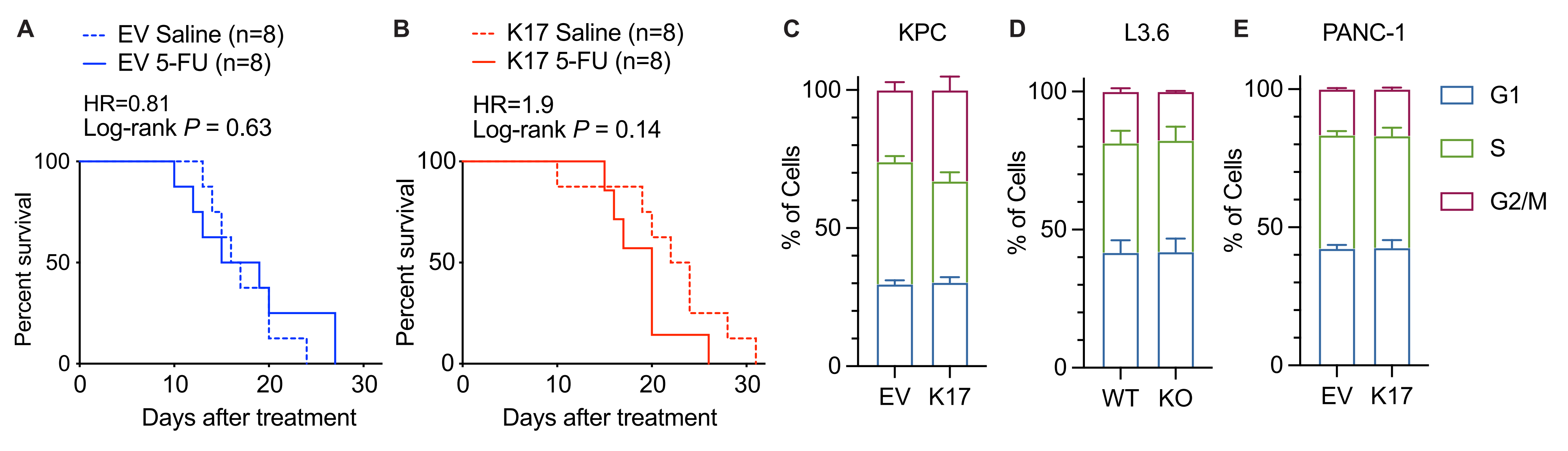
